# Oxidative switch drives mitophagy defects in dopaminergic *parkin* mutant patient neurons

**DOI:** 10.1101/2020.05.29.115782

**Authors:** Aurelie Schwartzentruber, Camilla Boschian, Fernanda Martins Lopes, Monika A Myszczynska, Elizabeth J New, Julien Beyrath, Jan Smeitink, Laura Ferraiuolo, Heather Mortiboys

**Affiliations:** Sheffield Institute for Translational Neuroscience, University of Sheffield, United Kingdom; School of Chemistry, University of Sydney, Australia; Khondrion BV, Philips van Leydenlaan 15, 6525EX Nijmegen, The Netherlands

## Abstract

Background Mutations in parkin are the most common cause of early onset Parkinson’s disease. Parkin is an E3 ubiquitin ligase, functioning in mitophagy. Mitochondrial abnormalities are present in parkin mutant models. Patient derived neurons are a promising model in which to study pathogenic mechanisms and therapeutic targets. Here we generate induced neuronal progenitor cells from parkin mutant patient fibroblasts with a high dopaminergic neuron yield. We reveal changing mitochondrial phenotypes as neurons undergo a metabolic switch during differentiation. Methods Fibroblasts from 4 controls and 4 parkin mutant patients were transformed into induced neuronal progenitor cells and subsequently differentiated into dopaminergic neurons. Mitochondrial morphology, function and mitophagy were evaluated using live cell fluorescent imaging, cellular ATP and reactive oxygen species production quantification. Results Direct conversion of control and parkin mutant patient fibroblasts results in induced neuronal progenitor and their differentiation yields high percentage of dopaminergic neurons. We were able to observe changing mitochondrial phenotypes as neurons undergo a metabolic switch during differentiation. Our results show that when pre-neurons are glycolytic early in differentiation mitophagy is unimpaired by PRKN deficiency. However as neurons become oxidative phosphorylation dependent, mitophagy is severely impaired in the PRKN mutant patient neurons. These changes correlate with changes in mitochondrial function and morphology; resulting in lower neuron yield and altered neuronal morphology. Conclusions Induced neuronal progenitor cell conversion can produce a high yield of dopaminergic neurons. The mitochondrial phenotype, including mitophagy status, is highly dependent on the metabolic status of the cell. Only when neurons are oxidative phosphorylation reliant the extent of mitochondrial abnormalities are identified. These data provide insight into cell specific effects of PRKN mutations, in particular in relation to mitophagy dependent disease phenotypes and provide avenues for alternative therapeutic approaches.

## Introduction

Parkinson’s disease (PD) is the second most common neurodegenerative disease, with approximately 10 million people affected worldwide. Only symptomatic treatment options are available. Mutations in *PRKN* are the most common cause of early onset PD (EOPD). Parkin is an E3 ubiquitin ligase and functions in the mitophagy pathway ^1^. Mitochondrial dysfunction is well established in both familial and sporadic forms of PD (recently reviewed ^2^). Mitochondrial abnormalities are present in both *PRKN* null Drosophila ^3^ and mice ^4^. We and others have shown mitochondrial abnormalities in peripheral cells from patients with *PRKN* mutations ^5–8^; these include cellular ATP defects, mitochondrial membrane potential deficiencies, complex I defect and altered mitochondrial morphology. Recent work suggests mitophagy is defective across many PD types ^9^. Several reports have found alterations in the same mitochondrial parameters in iPSC derived *PRKN* deficient neurons ^10–14^. These studies provide insight into a mitochondrial phenotype in *PRKN* deficient neurons. DA neurons are particularly vulnerable to mitochondrial abnormalities due to their high basal oxidative load, tonic activity and highly complex arborisation of the dendritic network ^15^. The studies so far have utilised the iPSC reprogramming route and subsequent differentiation into DA neurons, which generates a relatively poor yield of DA neurons. Therefore, the specific role and importance of mitochondrial abnormalities in a *PRKN* deficient background in DA neurons remains unclear.

The role of parkin in the mitophagy pathway is extremely well documented mainly in tumour cell lines over expressing parkin with mitophagy induction due to treatment with uncoupling agents such as CCCP. Limited studies have investigated the role of parkin dependent mitophagy in cells expressing endogenous parkin and fewer still without the induction of mitophagy via uncoupling. Recent findings from several *in vivo* models have called into question the relative importance of parkin dependent mitophagy in adult DA neurons with studies in PINK1 deficient mice and *PRKN* and PINK1 deficient Drosophila showing no difference in mitophagy rates in DA neurons ^16,17^. However an age dependent increase in mitophagy in DA neurons which is absent in *PRKN* and PINK1 deficient Drosophila has also been identified ^18^. Therefore more studies are needed to elucidate the importance of mitophagy in *PRKN* deficient EOPD.

Our study uses a direct conversion route from patient fibroblasts to induced neuronal progenitor cells (iNPC’s) and subsequently to DA neurons with a high yield. Direct reprogramming methods result in cells that both retain the genetic background and the age phenotype of the donor fibroblasts ^19^. With a high yield of DA neurons we are able to study the mitochondrial and mitophagy phenotypes throughout differentiation in this specific cell population. We show that mitophagy defects are dependent on the metabolic status of the cell; with high mitophagy rates in *PRKN* deficient neurons early in differentiation when the cells are mainly glycolytic and mitophagy rates which are extremely low in *PRKN* deficient neurons reliant upon oxidative phosphorylation. We also show treatment with known potent intracellular redox-modulating agents improves the neuronal phenotype of the neurons without restoring mitochondrial function or morphology.

## Results

### iNPC derived DA neurons display high yield and purity

Unlike iPSC reprogramming of fibroblasts, the direct conversion method first developed by Meyer et al produces iNPC’s which are tripotent and able to differentiate into neurons, astrocytes and oligodendrocytes ^20^. The iNPC’s produced using this methodology can be differentiated into specific neuronal populations with a high purity yield, as demonstrated for motor neurons ^21^. We successfully converted fibroblasts from 4 control and 4 *PRKN* mutant patients producing iNPC’s which displayed a clear change in cell morphology, proliferation rate and stained positive for the NPC markers, Pax6 and nestin as found previously for all iNPC’s reprogrammed using this methodology ^20–23^ (data not shown).

In order to be able to investigate biochemical parameters in DA neurons without other neuronal types or non-neuronal cells contaminating the culture; a high yield of DA neurons is required. We optimised the DA differentiation protocol based upon Swistowski and co-workers ^24^. Differentiation is in three stages, first iNPC’s are treated with DAPT, a Notch inhibitor that enhances neuronal differentiation; in stage 2 the cells are driven towards a rostral midbrain neuronal lineage and finally in stage 3 DA neuron differentiation is complete. As differentiation proceeds the cellular morphology alters; at day 15 of differentiation cells have become elongated and begun to form connections as compared to iNPC morphology and by day 27 the cells have formed longer, larger connections (Figure 1A shows brightfield images of cells at two stages of differentiation). The morphology is distinct from the parental iNPC morphology. However, we note, the processes are shortened and thicker than those usually seen in neurons differentiated from iPSC’s or primary embryonic cultures. Therefore, we sought to characterise the expression of several pan-neuronal markers and DA specific markers throughout differentiation. We assessed expression of pan neuronal markers βIII tubulin, MAP2 and NeuN at various stages of differentiation (day 12, day 17 and day 27). The amount of cells staining positive for each of these markers increases throughout differentiation, resulting at day 27 in 94.5% of cells staining for βIII tubulin, 87.9% for MAP2 and 76.5% for NeuN (quantification throughout differentiation is shown in Figure 1B-D with representative images at day 27 shown for each marker in a control and a *PRKN* mutant line). Next, we investigated the expression of two DA markers, tyrosine hydroxylase (TH) and the dopamine transporter (DAT). Again, expression increases throughout differentiation; with undetectable levels of TH and DAT expression at day 17 of differentiation. At day 27 of differentiation however 89.9% of cells stain positive for TH and 82.4% for DAT (Figure 1E and F quantification throughout differentiation and representative images at day 27 of differentiation). In addition, we quantified mRNA transcript level for TH at day 27 as compared to iNPC’s and found a 4.2 fold increase in DA neurons compared to iNPC’s (Figure 1G). Neurosensor 521 dye labels both noradrenaline and dopamine in live cells, and we found that at end stage of differentiation 87.8% of cells stained positive with Neurosensor 521 (Figure 1H). We next sought to measure the dopamine content and stimulated release of dopamine from our neuronal cultures. We found that the neurons had measurable intracellular dopamine and the neurons could be stimulated to release dopamine using potassium chloride in a magnesium containing buffer. *PRKN* mutant neurons contain less intracellular dopamine than controls (mean +/- SD, controls 111.8 +/- 2.6, parkin mutants 88.4 +/- 3.4 nmol/l dopamine; p < 0.001; Figure 1I). Upon stimulation with potassium chloride the amount of dopamine in the media increases in controls by 51% and only by 15% in *PRKN* mutant patients (Figure 1F). The neurons do not release dopamine when stimulated with potassium chloride without magnesium present. Finally, we also assessed the cellular membrane potential to assess neuronal properties of our cells in culture. We find positive staining for membrane potential and fluctuations as expected in active neuronal cultures (Figure 1J); however, the *PRKN* mutant neurons are less responsive to stimuli.

**Figure 1.**
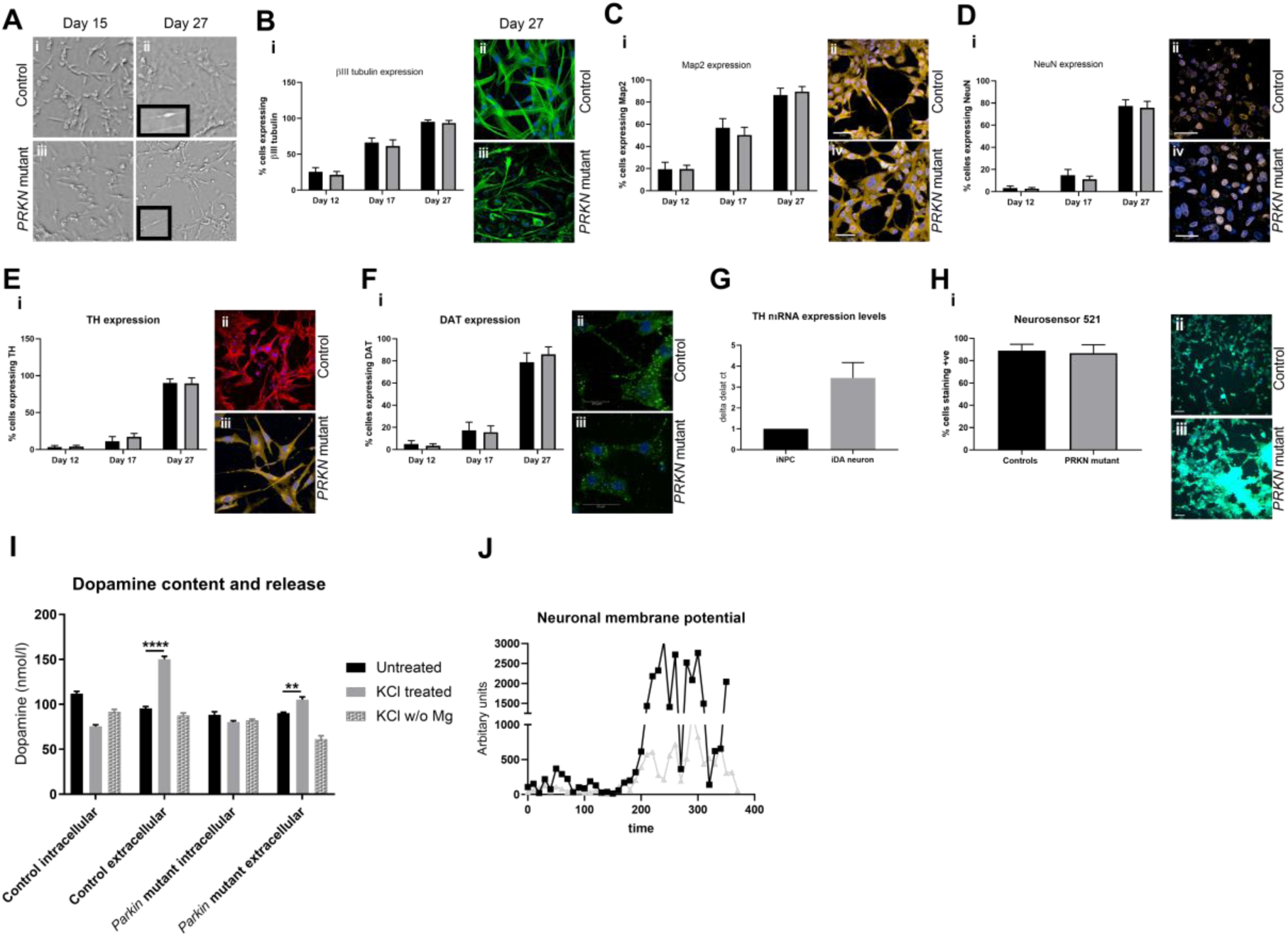
Control and *PRKN* mutant iNPC derived dopaminergic iNeurons characterization throughout differentiation. **A** shows bright field images (scale bar = 100µm) at day 15 and day 27. The insets show magnified regions highlighting the processes of the neurons. B-F shows (i) quantification of each neuronal marker throughout differentiation in controls (black bars) and *PRKN* mutants (grey bars) and (ii) shows representative images of a control and *PRKN* mutant at day 27. **B(ii)** green is βIII tubulin and blue nucleus. **C(ii)** red is MAP2 and blue the nucleus. **D(ii)** red is NeuN and blue nucleus. **E(ii)** red is tyrsosine hydroxylase and blue the nucleus. **F(ii)** green is DAT and blue the nucleus. **G** shows the quantification of mRNA expression levels for tyrosine hydroxylase in iNPC’s and day 27 neurons, showing a 4 fold increase in expression (black bars controls and grey bars *PRKN* mutants). **H(i)** shows the quantification at day 27 of Neurosensor staining which stains dopamine and noradrenaline showing 90% of cells staining positive in both controls (black bars) and *PRKN* mutants (grey bars). (ii) shows representative images of control and *PRKN* mutant at day 27. **I** shows the dopamine content and release assay in control and *PRKN* patient neurons. Intracellular and extracellular dopamine content assessed in neurons from three rounds of differentiation for each condition. Neurons are either untreated, stimulated with potassium chloride or potassium chloride without magnesium in the buffer. Extracellular dopamine levels increase when neurons are stimulated with potassium chloride in both control (**** p < 0.001) and *PRKN* mutant patient neurons (** p = 0.0032). **J** shows neuronal membrane potential in control (black line) and *PRKN* mutants (grey line) recorded at baseline and after stimulation at time 200 seconds. All quantification was done on at least three different differentiations of four control and four *PRKN* mutant neurons; two way ANOVA with Tukey multiple comparisons correction was used.

### *PRKN* mutant DA neurons display increased cell death and morphological abnormalities

Others have previously reported fewer surviving neurons at the end of differentiation towards a DA enriched population from iPSC’s derived from *PRKN* mutant parental patient fibroblasts ^14^. We investigated this during the differentiation from iNPC’s to DA neurons. There was significant cell death occurring throughout differentiation specifically in the *PRKN* mutant patient derived DA neurons; the percentage of cells surviving until the end of the differentiation was significantly reduced (mean +/- SD, controls 83.62 +/- 4.8; parkin mutants 52.72 +/- 11.98); however, the same % yield of surviving neurons expressed DA markers between controls and *PRKN* mutants. We quantified cell death using activated caspase 3 staining. The number of activated caspase 3 positive spots was higher in *PRKN* mutant neurons compared to controls at day 17, with a subsequent dramatic increase during the final stage of differentiation (% cells with activated caspase 3 staining at day 17 15% and at day 27 64%); whereas in control neurons the level remains constant at approximately 7.8% (Figure 2A). Furthermore, DA neurons from *PRKN* mutants displayed altered neuronal morphology at end stage differentiation; being more round and less elongated (Figure 2B and C; controls 1.92 +/- 0.02; *PRKN* mutants 2.74 +/- 0.2; p < 0.05 for cell roundness and controls 0.013 +/- 0.0002; *PRKN* mutants 0.008 +/- 0.001; p < 0.05 for cell elongation).

**Figure 2.**
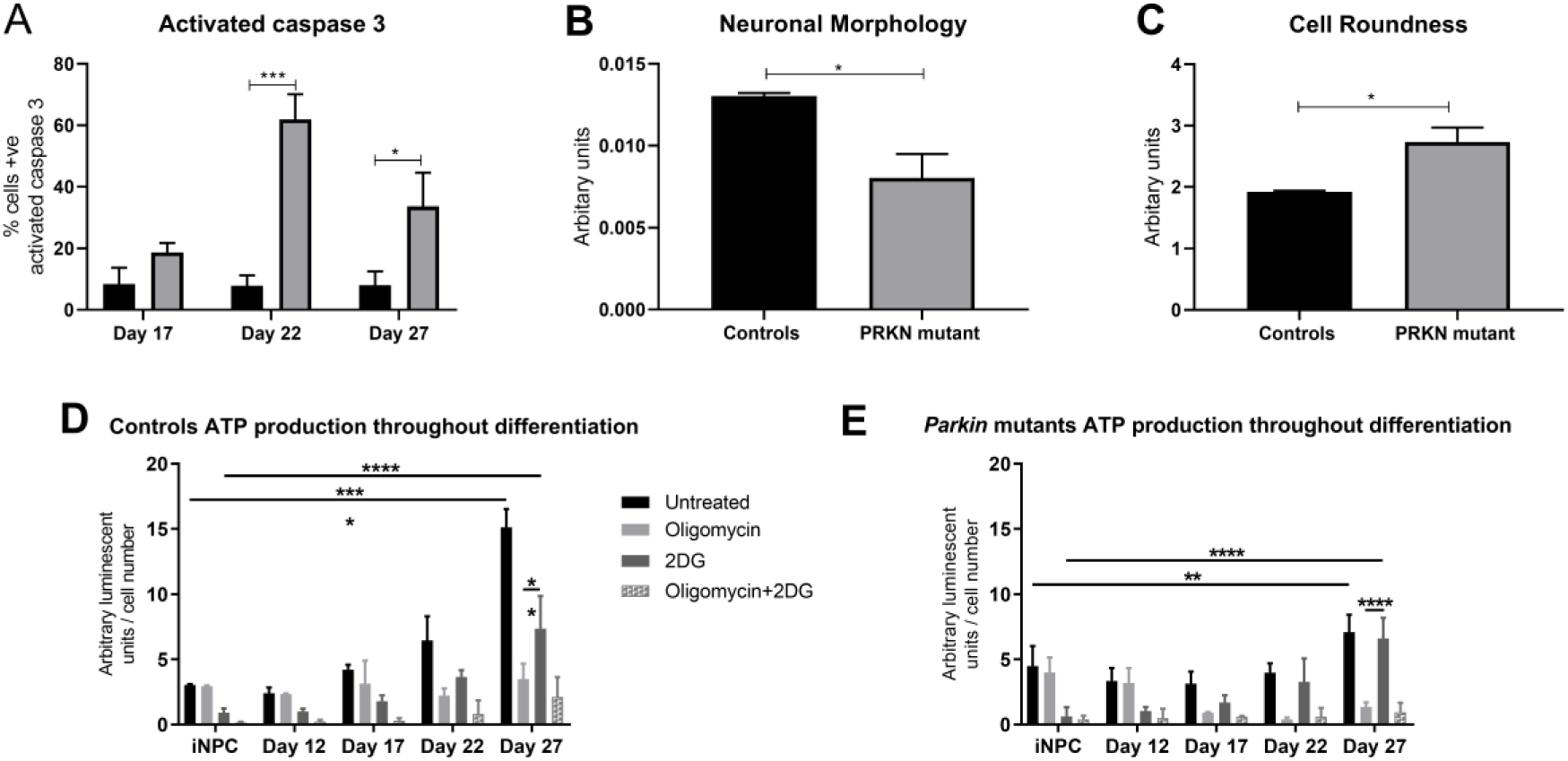
Neuronal morphology and metabolic status during differentiation. A Quantification of activated caspase 3 in controls (black bars) and *PRKN* mutants (grey bars) throughout differentiation. Significant increase in activated caspase 3 positive cells in *PRKN* mutants at day 22 (*** p = 0.0008) and at day 27 (* p = 0.036). **B** is a measure of neuronal branching; the *PRKN* mutant patient neurons have reduced neuronal branching compared to controls (* p = 0.029). **C** is a measure of the roundness of the cells. The *PRKN* mutant patient neurons are more round than control neurons (* p = 0.029). D **and E** show ATP production dependent on glycolysis or oxidative phosphorylation throughout differentiation (D shows control neurons and E shows *PRKN* mutant neurons). ATP levels increase throughout differentiation for both control and *PRKN* mutant neurons. iNPC’s rely wholly on glycolysis for ATP generation; this switches throughout differentiation until at day 27 neurons are ∼85% dependent on oxidative phosphorylation for ATP production (iNPC total ATP vs day 27 total ATP p = 0.0001; iNPC 2DG ATP vs day 27 2DG ATP p = 0.0001; day 27 oligomycin ATP vs day 27 2DG ATP p = 0.005). This switch happens in both control and *PRKN* mutant neurons (iNPC ATP vs day 27 ATP p = 0.003; iNPC 2DG ATP vs day 27 2DG ATP p = 0.0001; day 27 2DG ATP vs day 27 oligomycin ATP p = 0.0001. All experiments were repeated on three separate rounds of differentiation in each control and *PRKN* mutant patient line (four different controls and *PRKN* mutant patient lines are included). Bar graphs represent mean with SD. All statistics done by two-way ANOVA test using Sidaks multiple comparisons test.

### Metabolism shifts throughout differentiation; revealing mitochondrial morphology and functional abnormalities in *PRKN* mutant DA neurons

Recent work has shown direct reprogramming methods retain the age characteristics of the donor fibroblasts; importantly this also includes the switch to oxidative phosphorylation during direct reprogramming and reductions in mitochondrial function and gene expression in the neurons generated from aged donors rather than those generated from younger donors^19^. We therefore sought to understand the metabolism in our system, which utilises direct conversion of fibroblasts to iNPCs rather than reprogramming to iPSCs. We assessed the contribution of glycolysis and oxidative phosphorylation to the ATP levels in the cells from parental fibroblasts, to iNPC’s and at various stages throughout neuronal differentiation. Both the parental fibroblasts and the iNPC’s are wholly glycolytic with inhibition of complex V of the respiratory chain resulting in no decrease in energy levels in the cells (Figure 2D; untreated 3.1 +/- 0.6; OXPHOS inhibited 2.9 +/- 0.7; glycolysis inhibited 0.9 +/- 0.3). However during neuronal differentiation the cells undergo a metabolic switch from glycolysis to oxidative phosphorylation; such that by the end stage of differentiation neurons are reliant on oxidative phosphorylation for 88% of their energy generation (Figure 2D, untreated 15.1 +/- 1.4; OXPHOS inhibited 3.5 +/- 1.2; glycolysis inhibited 8.5 +/- 2.5). This switch occurs in both control and *PRKN* mutant DA neurons (Figure 2D and E). It is interesting to observe the time point at which the dramatic increase in activated caspase 3 and cell death occurs in the *PRKN* mutant neurons (day 22) correlates with the metabolic switch towards OXPHOS reliance.

In order to fully understand mitochondrial function and morphology as this metabolic switch occurs and the role of parkin in this; we investigated mitochondrial function, morphology and mitophagy throughout differentiation in control and *PRKN* mutant neurons. Previous reports have shown mitochondrial fragmentation in *PRKN* mutant iPSC derived neurons ^25,26^. We observe the same mitochondrial fragmentation at the end stage of differentiation accompanied by an increase in mitochondrial number (Figure 3A mitochondrial interconnectivity: controls 0.07 +/- 0.003; *PRKN* mutants 0.04 +/- 0.005 p < 0.05; Figure 3B mitochondrial number (% normalised to controls): controls 100 +/- 3.4; *PRKN* mutants 204 +/- 35; p < 0.0001;). In both control and *PRKN* mutant DA neurons throughout differentiation mitochondria become more interconnected as the metabolic switch occurs. The controls then return to a ‘normal’ morphology once this has happened (Figure 3A). The increase in mitochondrial number in *PRKN* mutant DA neurons does not seem to be driven by increased biogenesis but rather the total mitochondrial content remains fairly constant however mitochondria are smaller and more fragmented in the *PRKN* mutant DA neurons. There is much debate in the literature as to whether the energy defect or increased ROS production is more detrimental in PD. We found dramatically increased mitochondrial ROS levels at end stage of differentiation (controls 0.17 +/- 0.018; *PRKN* mutants 0.6 +/- 0.15; * p < 0.05; Figure 3C). There is no change in mitochondrial ROS levels at earlier stages of differentiation. In terms of mitochondrial function, we show that mitochondrial membrane potential is significantly reduced only at end stage differentiation (MMP controls 0.02 +/- 0.004, *PRKN* mutant 0.003 +/- 0.002, **** p < 0.001; Figure 3D). However, there is a worsening trend in MMP decreases as differentiation continues. A similar pattern is observed for cellular ATP levels with increasing deficits as differentiation progresses. The first significant decrease in ATP levels is observed when the neurons are becoming reliant on oxidative phosphorylation at day 17 of differentiation (controls 100 +/- 5.5, parkin mutants 82 +/- 4.3, * p < 0.05; Figure 3E). At the end stage of differentiation the deficit is more severe (% normalised to controls: controls 100 +/- 15, *PRKN* mutant 61 +/- 15; **** p < 0.01; Figure 3E). Taking the above data together we see dramatic changes in mitochondrial function and morphology, which are only revealed in *PRKN* mutant neurons as the metabolic switch occurs from glycolysis to oxidative phosphorylation; accompanied by a dramatic increase in mitochondrial ROS levels once this switch has taken place.

**Figure 3.**
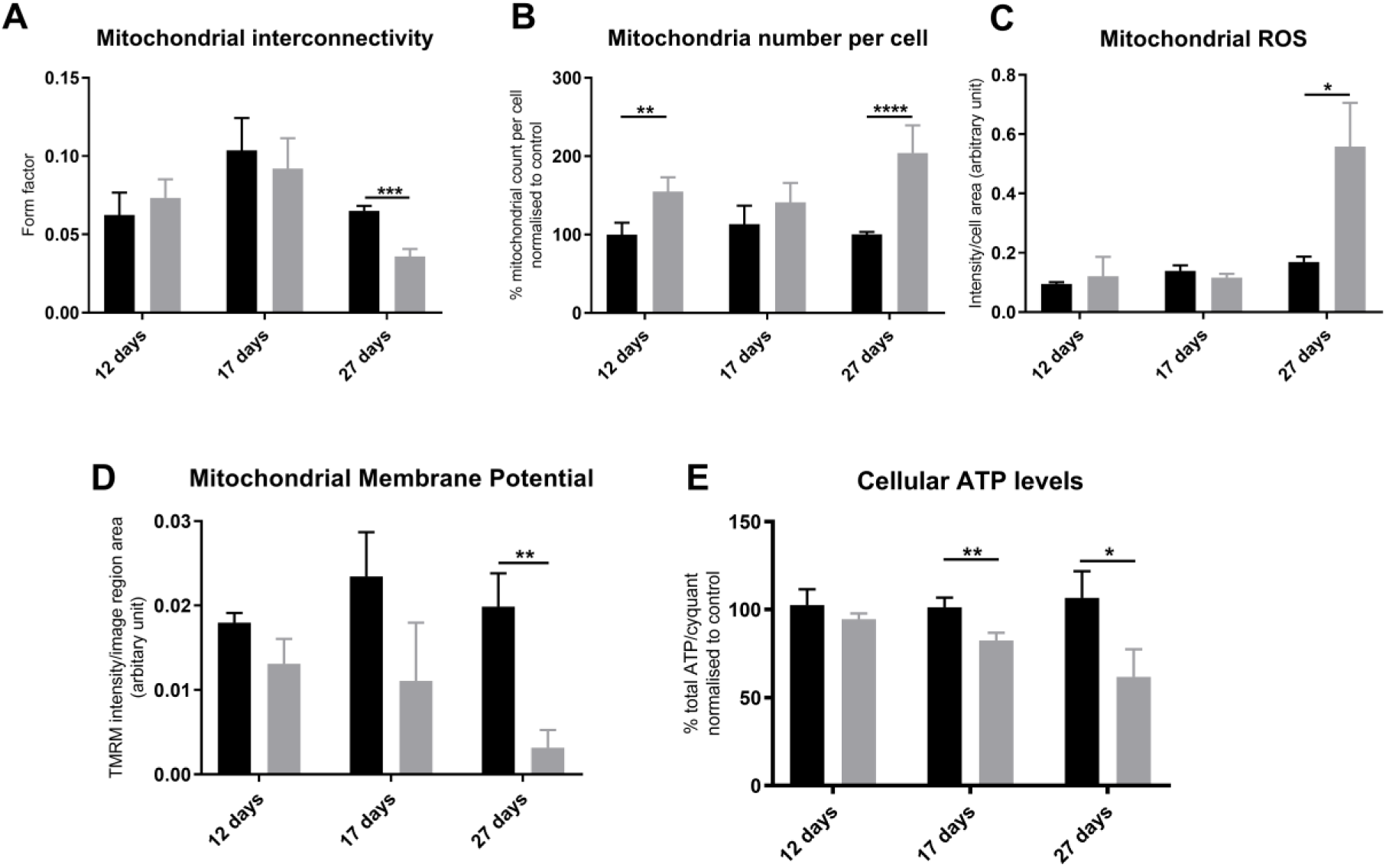
Mitochondrial morphology and function in dopaminergic iNeurons throughout differentiation. A shows mitochondria in both control (black bars) and *PRKN* mutant patient neurons (grey bars) elongate as the metabolic switch from glycolysis to oxidative phosphorylation begins at day 17 of differentiation. By day 27 of differentiation mitochondria in *PRKN* mutant patient neurons are fragmented compared to control neurons (*** p = 0.0004). **B** shows mitochondrial number is increased in *PRKN* mutant neurons (grey bars) compared to controls (black bars); (at day 12 ** p = 0.008 and day 27 **** p < 0.0001). **C** shows mitochondrial ROS production is significantly increased at day 27 of differentiation in *PRKN* mutant neurons (grey bars) compared to controls (black bars; * p = 0.037). **D** shows mitochondrial membrane potential is decreased in *PRKN* mutant patient neurons (grey bars) throughout differentiation, however the most dramatic and only significant reduction is at day 27 of differentiation (** p = 0.003). **E** shows cellular ATP levels are reduced in *PRKN* mutant patient neurons at day 17 (** p = 0.005) and day 27 (* p = 0.01) of differentiation. All experiments were repeated on three separate rounds of differentiation in each control and *PRKN* mutant patient line (four different controls and *PRKN* mutant patient lines are included). Bar graphs represent mean with SD. All statistics done by two-way ANOVA test using Sidaks multiple comparisons test.

### Mitophagy rates are higher in *PRKN* mutant neurons with glycolytic capacity before becoming defective when neurons undergo a metabolic switch

As parkin is known to function in a well characterised parkin dependent mitophagy pathway targeting dysfunctional or damaged mitochondria for degradation; we developed a live imaging assay to assess both basal and induced mitophagy rates in the neurons throughout differentiation. This assay relies upon live staining of the total mitochondrial population and the lysosomal population combined with advanced high content imaging acquisition, data processing and analysis. At 12 days differentiation, when cells are positive for the pan neuronal marker βIII tubulin but are not yet DA and are glycolytic, *PRKN* mutant neurons have very similar rates of basal mitophagy as controls (controls 0.08 +/- 0.4; *PRKN* mutants 0.09 +/- 0.6, Figure 4Ai). When mitophagy is induced in these neurons the *PRKN* mutant neurons mount a higher response to global mitochondrial inhibition but they cannot sustain mitophagy for as long as the control neurons are able to (Figure 4Aii).

**Figure 4.**
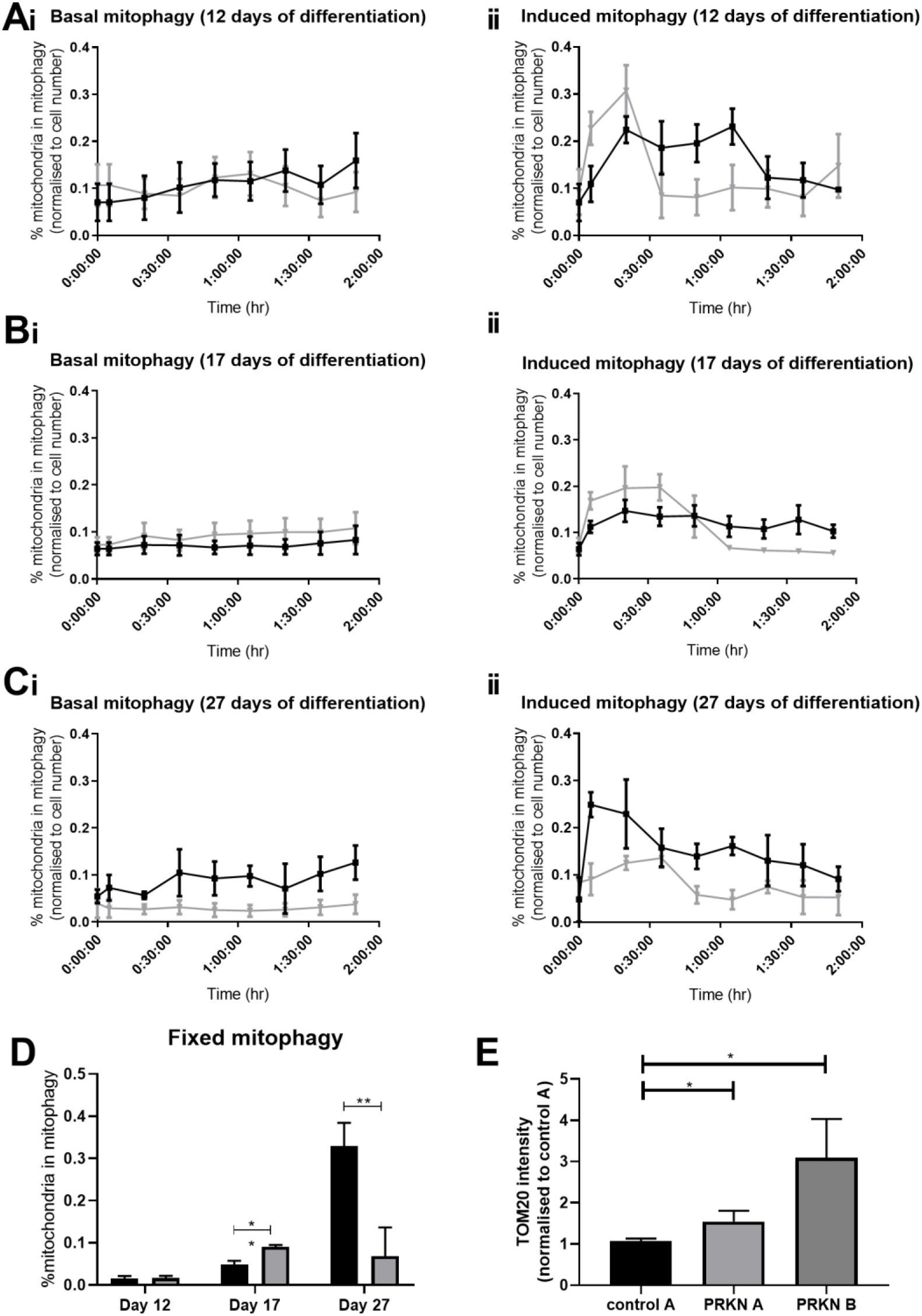
Mitophagy throughout dopaminergic iNeuron differentiation. **A, B and C** Quantification of percentage of mitochondria undergoing basal (i) or induced (ii) mitophagy per cell over time at day 12 (**A**), day 17 (**B**) or day 27 of differentiation (**C**). Graphs represent the quantification of mitochondria undergoing mitophagy over time. Basal mitophagy is unaltered in *PRKN* mutants (grey lines) compared to controls (black lines) at day 12 of differentiation (Ai); however by day 17 basal mitophagy is increased in *PRKN* mutants (p = 0.019; Bi) and at endpoint of differentiation day 27, *PRKN* mutants have significantly reduced basal mitophagy levels (* p = 0.011, Ci). Induced mitophagy is increased in *PRKN* mutants initially after induction at day 12 (Aii); the same pattern is seen at day 17 (Bii) however by day 27 induced mitophagy is significantly lower in *PRKN* mutant neurons (* p = 0.0113, Cii). **D** shows quantification of mitophagy using an alternative measure; showing the same pattern as the live assay. No difference at day 12, an increase at day 17 in the *PRKN* mutants (grey bars) compared to controls (black bars, ** p = 0.0016) and a reduction at day 27 (** p = 0.0035). All experiments were repeated on three separate rounds of differentiation in each control and *PRKN* mutant patient line (four different controls and *PRKN* mutant patient lines are included). Bar graphs represent mean with SD. **E** shows quantification of the amount of Tom20. *PRKN* mutant neurons have increased Tom20 amounts at day 27 of differentiation as compared to control A (*PRKN* A * p = 0.04, *PRKN* B * p = 0.02). n = 3 for each line.

However, at day 17 of differentiation when neurons are approximately 50/50 reliant on glycolysis and oxidative phosphorylation *PRKN* mutant neurons have higher basal mitophagy levels than controls (controls 0.07 +/- 0.03, *PRKN* mutants 0.09 +/- 0.05, Figure 4Bi) and again mount a higher response to mitochondrial inhibition but cannot sustain that level of mitophagy overtime (Figure 4Bii). Finally, at the end stage of differentiation when neurons are reliant on oxidative phosphorylation *PRKN* mutant neurons have a severe deficit in basal and induced mitophagy (Figure 4C basal mitophagy controls 0.09 +/- 0.025, *PRKN* mutants 0.025 +/- 0.008; induced mitophagy controls 0.3 +/- 0.09, *PRKN* mutants 0.1 +/- 0.01, p < 0.01). We used two alternative methods of evaluating mitophagy rates previously validated ^27,28^; using these method we found very similar results throughout differentiation and at endpoint (Figure 4D and 4E). Tom20 amount is increased in *PRKN* mutant patient derived neurons, indicating less mitophagy aligning with the mitophagy rates measured using the live and fixed assays.

### Redox modulating compounds KH176 and KH176m partially reverses neuronal deficits in *PRKN* mutant DA neurons

In order to evaluate if the driving mechanism in *PRKN* mutant DA neurons is the loss of energy production by the mitochondria or the dramatically increased mitochondrial ROS production, we treated the neurons with the known potent intracellular redox-modulating agents KH176 and KH176m ^29^ currently in clinical trials in mitochondrial patients with m.3243A>G spectrum disorders ^30^. In agreement with the mechanism of action^28^ treatment with KH176 and KH176m decreased mitochondrial ROS production to control levels after 24 hours treatment (Figure 5A, control A vehicle treated 0.16 +/- 0.006; control A KH176 treated 1µM 0.14 +/- 0.02; *PRKN* mutant A vehicle treated 0.7 +/- 0.15; *PRKN* mutant A KH176 treated 1µM 0.12 +/- 0.05; p < 0.001). KH176 and KH176m treatment had no significant effects on mitochondrial function (MMP or cellular ATP levels) or mitochondrial morphology parameters (Figure 5 B-D). Treatment with KH176 and KH176m did show a mild effect on neuronal morphology after only 24 hours of treatment; with the resulting neurons being less round after treatment; a morphology closer to control neurons (Figure 5E, cell roundness: control A vehicle treated 1.5 +/- 0.09; control A KH176 treated 1µM 1.6 +/- 0.06; *PRKN* mutant A vehicle treated 3.1 +/- 0.6; *PRKN* mutant A KH176 treated 1µM 2.2 +/- 0.6). KH176 has not been assessed before for an effect on mitophagy rates; we investigated if KH176m treatment could affect basal mitophagy rates in neurons from *PRKN* mutant A. Our data show a significant increase in basal mitophagy rates after treatment with KH176m in both control and *PRKN* neurons (Figure 5F, control A vehicle treated 1.14 +/- 0.11, control A KH176m 100nM 1.27 +/- 0.11, control A KH176m 1000nM 1.56 +/- 0.15 *PRKN* mutant A vehicle treated 0.6 +/- 0.05, *PRKN* mutant A KH176m 100nM 0.9 +/- 0.05, *PRKN* mutant A KH176m 1000nM 0.8 +/- 0.08).

**Figure 5.**
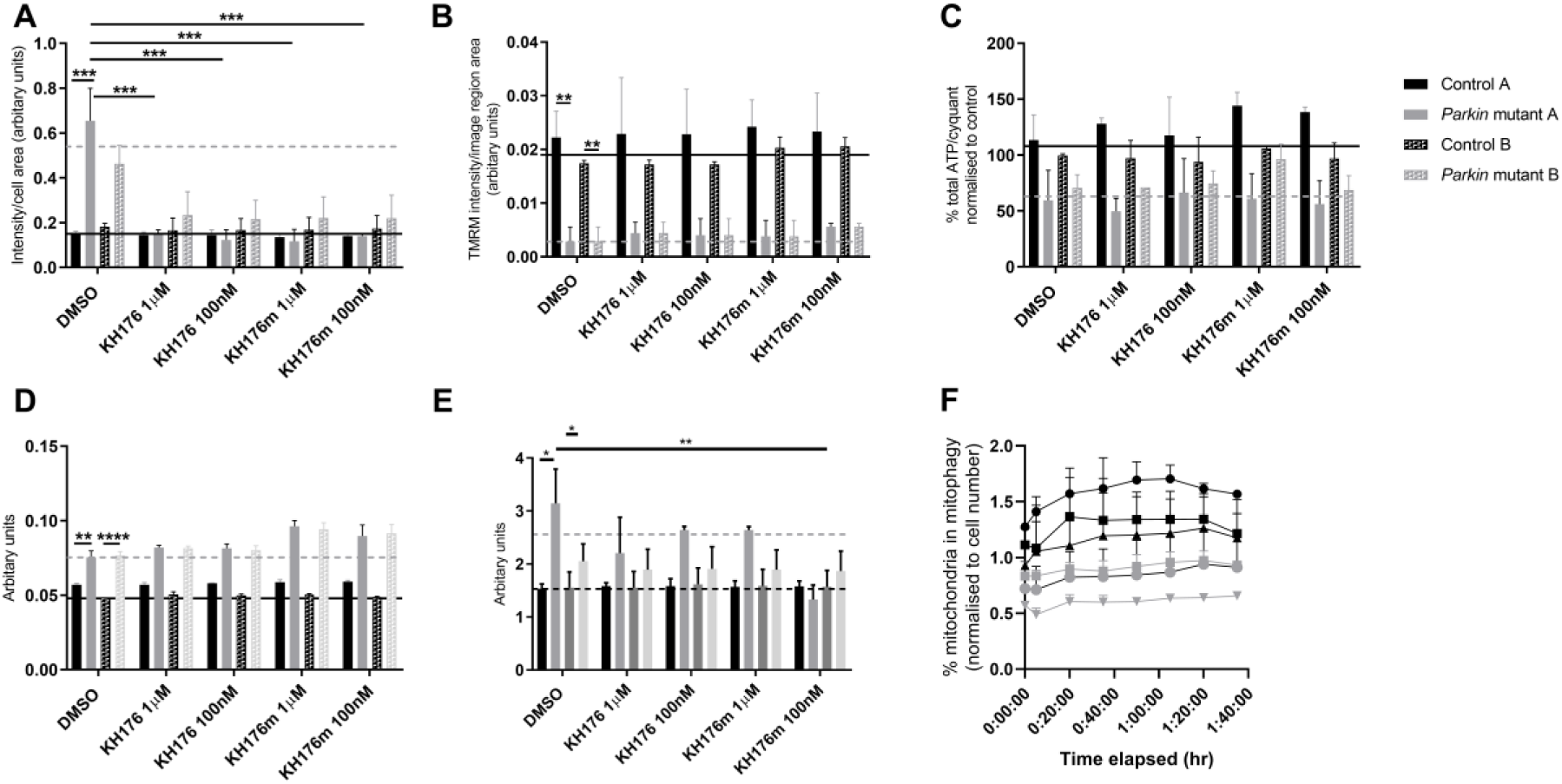
Treatment of dopaminergic iNeurons with KH176 and KH176m. **A** shows mitochondrial ROS levels are significantly reduced with treatment of both KH176 and KH176m (n = 3 for each line presented, 2 way ANOVA with Sidaks multiple comparisons test *** p < 0.0001). **B** shows mitochondrial membrane potential is significantly reduced in *PRKN* mutant patient neurons however treatment with KH176 and KH176m has no effect on mitochondrial membrane potential (** p = 0.0009 and 0.025 respectively). **C** shows cellular ATP levels are reduced in *PRKN* mutant neurons however treatment with KH176 and KH176m has no significant effect. **D** shows mitochondria are more round in *PRKN* mutant patient neurons compared to controls, again treatment with KH176 and KH176m has no effect (** p = 0.0048 and **** p = 0.0001 respectively). **E** shows neuronal roundness is increased in *PRKN* mutant neurons; treatment with KH176m at 100nM has a significant effect of reducing neuronal roundness, indicating the neurons are more elongated and similar in morphology to the controls (* p = 0.018 and ** p = 0.0045 respectively). For A-E the black dotted line shows mean vehicle treated for controls and grey dotted line mean vehicle treated for *PRKN*, n = 3 for each line. Two way ANOVA with Sidaks multiple comparisons test used. **F** Basal mitophagy is reduced in *PRKN* mutant A (grey triangles) compared to control A (black triangles); treatment with KH176m at both 100nM (squares) and 1000nM (circles) concentrations increase basal mitophagy rates in both control A and *PRKN* mutant A (n = 2 for each line presented, 2 way ANOVA with Sidaks multiple comparisons test control vehicle vs control KH176m 1000nM p = 0.0088; control vehicle vs *PRKN* vehicle p = 0.0003; *PRKN* mutant A vehicle vs *PRKN* mutant A KH176m 100nM p = 0.0001).

## Discussion

Our study is the first to report successful reprogramming via the iNPC route of PD *PRKN* mutant patient fibroblasts; varying reprogramming methods depend on competent energy generation for successful reprogramming^19^. We have previously reported severe mitochondrial abnormalities in *PRKN* mutant fibroblasts ^5^; reprogramming of fibroblasts with a reduction in metabolic function can be challenging using iPSC routes^19^ however here we show metabolically challenged fibroblasts can be reprogrammed using this direct reprogramming route. Recently others have used alternative direct reprogramming methods to generate dopamine like neuronal cells from sporadic and LRRK2 Parkinson’s patient cells^31–33^. These studies showed several alternative reprogramming routes can lead to viable dopaminergic neuronal like cells; with each group assessing the dopaminergic qualities of the cells produced. Furthermore we have recently reported use of this reprogramming route to generate dopaminergic neurons from sporadic PD fibroblasts; in that study we found the mitochondrial abnormalities exasperated in the neurons compared to the fibroblasts from the same patient^34^. The specific method we have used here, the iNPC derived route has proved a useful model to study familial and sporadic forms of neurodegenerative diseases thus far; astrocytes derived from Motor Neuron Disease (MND) patients display neuronal toxicity when in co-culture with WT neurons similar to that seen with primary astrocytes from post-mortem biopsies from MND patients. Furthermore both iAstrocytes and iNeurons were recently used to investigate mechanisms of cell toxicity in C9orf72 dependent Motor Neuron Disease^20–23,35,36^. We report the generation of high yield of DA neurons from iNPC’s. The DA yield we achieved via this method is higher than reported via the iPSC differentiation route and similar or higher than that found with alternative direct reprogramming routes. We also note, the processes of the neurons generated via iNPC reprogramming are shorter than those from iPSC derived neurons. We hypothesize this is due to the retention of age characteristics during direct reprogramming methods; whereas iPSC derived neurons are more embryonic in nature and similar to primary cultures generated from mice. However, this requires further investigation to fully understand the mechanisms involved. We find more cell death in the *PRKN* mutant DA neurons throughout differentiation; less efficient differentiation has been reported by others for *PRKN* mutant patient neurons via iPSC reprogramming route ^13^. We also report iNPC derived *PRKN* mutant DA neurons are smaller and less elongated than controls. The increased cell death has been suggested to be dependent on the mitochondrial status of the cell rather than genotype ^37^. Our data would support this however further work to systematically test this would be useful to assess correlation between genotype and metabolic status.

Our study investigating mitochondrial function and morphology throughout differentiation suggests the increased cell death seen in *PRKN* mutant DA neurons co-insides with the neurons undergoing a metabolic switch from glycolysis to oxidative phosphorylation. We show the iNPC derived neurons have a clear switch in metabolism at day 22 with dependence on oxidative phosphorylation rather than glycolysis with a concurrent increase in the total amount of ATP in the neurons. This allows us to study mitochondrial function in these neurons which are metabolically more aligned to adult neurons *in vivo* (which are oxidative phosphorylation dependent) rather than embryonic neurons (glycolysis dependent) ^38^. Our data show that, while this metabolic switch is occurring, mitochondrial morphology changes; as the neurons become more oxidative phosphorylation dependent the mitochondria become more interconnected in both controls and *PRKN* mutants; we suggest this a change in mitochondrial morphology to allow the neurons to become reliant on oxidative phosphorylation. Once the metabolic switch has occurred mitochondrial morphology can return to the normal shape; however *PRKN* mutant neurons once OXPHOS dependent have increased mitochondrial fragmentation. This is opposite to the mitochondrial morphology phenotype we have previously reported in *PRKN* mutant fibroblasts ^5^; however others in the literature have previously reported a more fragmented mitochondrial network associated with *PRKN* deficiency ^39^; this is likely to be a cell type specific effect; our data suggesting this is dependent on the metabolic status of the cells.

Previous studies utilising iPSC derived *PRKN* mutant neurons have found mitochondrial abnormalities including defective mitophagy when induced using CCCP ^12^; however recent *in vivo* data from mouse and *Drosophila* models have shown little reduction in mitophagy on a *PRKN* or PINK1 deficient background ^16^. Here we show basal and induced mitophagy levels in *PRKN* mutant patient derived DA neurons; furthermore we find in a *PRKN* mutant background mitophagy levels are dependent on cellular energetic status. In cells which are dependent on glycolysis for energy production, basal and induced mitophagy are increased (or at least the same as controls) in *PRKN* mutant patient cells however upon the switch to OXPHOS dependency the *PRKN* mutant DA neurons have impaired basal mitophagy and are unable to mount a response to global mitochondrial dysfunction. Our data support the finding in *PRKN* deficient *Drosophila* that adult neurons increase levels of mitophagy during ageing however *PRKN* deficient neurons cannot ^18^. The specific mitophagy pathway being utilised in these *PRKN* mutant neurons is not clear and requires further investigation.

Although mitochondrial abnormalities have been clearly identified by many in PD models; there is debate as to whether the detrimental component of this is actually loss of energy or increased ROS production. Here we show that mitochondrial ROS levels are significantly increased only at end stage of differentiation when the neurons are OXPHOS dependent and have severe mitochondrial abnormalities. The increase in mitochondrial ROS is striking in all four *PRKN* mutant patient neuron lines. Previous studies have shown an increase in ROS in some *PRKN* mutant patient neurons but not in others and have measured total cellular ROS rather than mitochondrial specific ROS which could explain why the data we present here is more consistent across the group of patients. Targeting mitochondrial dysfunction for a potential therapeutic to slow or stop disease progression is an attractive option with many mitochondrial targeted therapeutics shown to be effective in various models of PD (recently reviewed ^2,40^). Different therapeutic strategies are being developed; some primarily acting to boost energy deficits whilst others are targeting ROS production. Here we show that treatment with the known redox-modulating compounds KH176 and KH176m dramatically reduces the mitochondrial ROS production with no significant effect on MMP or cellular ATP levels; however, KH176 and KH176m do have a mild beneficial effect on the neuronal morphology of the *PRKN* mutant neurons. These effects could be modulated by an increase in basal mitophagy after treatment with KH176m. This suggests a reversal of the energy deficit may not be required to have beneficial neuronal effects; however further work need to be done to fully investigate this, particularly over a longer term treatment.

In conclusion, our study utilises the iNPC technology to generate a high DA population of neurons which both express markers of DA neurons and release dopamine upon induction. Our data shows a predominant mitochondrial dysfunction present in these neurons which is far more pronounced than that found in the primary patient fibroblasts^5^. Our study builds on previous work as for the first time neuronal properties, mitochondrial functional, morphological and mitophagy parameters are assessed in the same neurons; neurons which all express TH and contain dopamine. Finally, our study highlights mitophagy as an energetic dependent process, which, in a *PRKN* mutant background varies considerably if the cells are glycolytic or OXPHOS dependent. This underlines the need to study mitophagy processes with endogenous levels of proteins in cell types which are relevant for disease and understand the energetic profile of the cells in order to be able to relate the findings to disease mechanism. Further studies to undertake detailed biochemical assessments of neuronal metabolism in this model in addition to utilising this model to assess putative neuroprotective compounds are warranted.

## Methods

### Culture of primary fibroblasts, generation and culture of iNPC’s

Primary fibroblasts were obtained from Coriell Cell Repository (coriell.org) controls: ND29510, GM09400, GM23967 and AG06882; *PRKN* mutant patients: ND30171, ND31618, ND40067 and ND40078 (details of mutation are given at coriell.org; full information is now available from NINDS data repository). Control and *PRKN* mutant groups were age and sex matched (controls 57 +/- 6.8; parkin mutants 53 +/- 8 years). Fibroblasts were cultured in EMEM as previously described ^5^. iNPC’s were generated as previously described ^41^. iNPC’s were maintained in DMEM/Ham F12 (Invitrogen); N2, B27 supplements (Invitrogen) and FGFb (Peprotech) in fibronectin (Millipore) coated tissue culture dishes and routinely sub-cultured every 2-3 days using accutase (Sigma) to detach them.

### Neuron differentiation of iNPC’s

Neurons were differentiated from iNPC’s as previously described^34^. Briefly, iNPCs are plated in a 6- well plate and cultured for 2 days in DMEM/F-12 medium with Glutamax supplemented with 1% NEAA, 2% B27 (Gibco) and 2.5µM of DAPT (Tocris). On day 3, DAPT is removed and the medium is supplemented with 1µM smoothened agonist (SAG; Millipore) and FGF8 (75ng/ml; Peprotech) for additional 10 days. Neurons are replated at this stage. Subsequently SAG and FGF8 are withdrawn and replaced with BDNF (30 ng/ml; Peprotech), GDNF (30 ng/ml; Peprotech), TGF-b3 (2 mM; Peprotech) and dcAMP (2 mM, Sigma) for 15 days.

### Immunofluorescence staining, live fluorescent imaging and ELISA

Neurons were plated and underwent immunocytochemistry staining as described previously^34^. Cells are plated into 96 well plates and fixed using 4% paraformaldehyde for 30 minutes. After PBS washes cells are permeabilised using 0.1% Triton X-100 for 10 minutes and blocked using 5% goat serum for 1 hour. Cells are incubated with primary antibodies (Pax6 (Abcam); nestin (Abcam); GFAP (Abcam), tyrosine hydroxylase (Abcam); DAT (ThermoFisher); β III tubulin (Millipore); Tom20 (BD Biosciences); LC3 (MBL); activated caspase 3 (Cell Signalling); Map2 (Abcam); NeuN (Abcam)); at 4 degrees for 16 hours. Cells are washed using PBS-Tween and incubated with Alexa Fluor conjugated secondary antibodies 488 and 568 (Invitrogen) and Hoescht (Sigma) 1µM prior to imaging. Imaging was performed using the Opera Phenix high content imaging system (Perkin Elmer). Twenty fields of view were imaged per well; in seven z planes. Images were analysed using Harmony software; maximum projections were used for analysis.

Dopamine ELISA was performed as per the manufacturer’s instructions (Labor Diagnostika Nord GmbH&Co. KG). Dopamine release experiments, neurons were incubated in HBSS with Ca^2+^ and Mg^2+^ (Gibco by Life Technologies) for 30minutes, or HBSS with Ca^2+^ and Mg^2+^ for 15 minutes and 56mM KCl (Fisher chemical) for another 15 minutes or HBSS without Ca^2+^ and Mg^2+^ (Gibco by Life Technologies) with 2mM EDTA for 15min and then 56mM KCL is added for another 15 minutes. Media is collected immediately; cells are harvested using accutase, centrifuged at 400g for 4min and resuspended in 10µl of PBS. EDTA 1mM and Sodium Metabisulfite (Sigma) 4mM are added to both the media and pellet to preserve the dopamine. The ELISA was read on a PheraStar plate reader (BMG Labtech) as per the manufacturer’s instructions; using the provided standard curve to calculate dopamine concentrations.

Neurons were incubated with 0.1 µM Neurosensor 521 (Sigma) and 1µM Hoechst in media for 30 minutes at 37 degrees. Cells were washed in phenol red free media and imaged using InCell 2000 (GE Healthcare) using 60x objective and 488nm excitation for Neurosensor 521 and 405nm excitation for Hoechst (method modified from ^42^). Fifteen fields of view per well were imaged and at 3 wells per line on at least three rounds of differentiation.

Neuronal membrane potential was measured using Fluovolt Membrane Potential Kit (ThermoFisher) as per the manufacturer’s instructions. Experiments were performed under basal or depolarizing conditions after treatment with isotonic potassium chloride solution (140 mM KCl, 5 mM NaCl, 1.8 mM CaCl2, 1.0 mM MgCl2, 20 mM HEPES, 20 mM Glucose, pH 7.4).

### Mitochondrial function, morphology and mitophagy measurements

Neurons were plated in 96 well plates; for MMP and morphology live imaging cells are incubated for one hour at 37 degree with 50nM tetramethylrhodamine (TMRM), 1µM rhodamine 123 and 1µM Hoescht (Sigma), after removal of dyes and replacement with phenol red free media plates are imaged using the Opera Phenix. Fifteen fields of view are imaged per well, in seven z planes. Images are analysed using Harmony software (Perkin Elmer). Segmentation protocols were established to segment the nuclei, mitochondria, and image region containing cytoplasm including projections. Analysis of number, size, intensity and morphology of mitochondria were calculated per image region using Harmony software (Perkin Elmer) using similar methodology as previously established ^27^. Cellular ATP measurements are undertaken using ATPLite kit (Perkin Elmer) as per manufacturer’s instructions. To assess dependency on OXPHOS or glycolysis, cells were pre-treated with oligomycin (Sigma) 10µM and 2-Deoxy Glucose (Sigma) 50mM for 30minutes at 37 degree and then ATP measurements were performed ^43^. Mitochondrial reactive oxygen species generation was assessed using mitochondrial NpFR2 probe ^44^ incubated with cells at 20µM and 1µM Hoechst for 30mins at 37°C, probes were removed and phenol red free media replaced. Cells were imaged using the Opera Phenix (Perkin Elmer). In order to assess mitophagy in live cells, cells were incubated for one hour at 37°C with 1µM tetramethylrhodamine (TMRM), 1µM Lysotracker Green (Invitrogen) and 1µM Hoescht, before washing to remove fluorescent probes. For the measurement of induced mitophagy 2µM Antimycin A (Sigma) and 5µM oligomycin (Sigma) were added prior to imaging. Images were captured in time lapse every 18 minutes in the same fields of view, minimum 6 fields of view per well. Images generated from the live imaging experiments were analysed using Harmony (Perkin Elmer software). We developed protocols in order to segment nucleus, image region containing cytoplasm, mitochondria, lysosomes, autolysosomes containing mitochondria. Maximal projection images were used for analysis. Mitochondria contained within lysosomes segmentation was set up in such a way to identify a mitophagy event when the overlap between mitochondria and lysosome was 100%.

Staining of cells using LC3 and Tom20 (as described above) and subsequent imaging using Opera Phenix and image analysis was used to validate the live imaging mitophagy assay. The image segmentation and analysis was set up in Harmony software as previously published ^27^. Furthermore as an additional mitophagy read out we utilised the loss of Tom20 signal from cells as previously determined ^28^.

### Statistical tests

All experiments were performed on at least triplicate differentiations for each control and *PRKN* mutant neuron or iNPC unless otherwise stated. Data are presented as mean +/- standard deviation. Students t test was used when comparing between control and *PRKN* mutant patients. When comparing different timepoints throughout differentiation a matched two way ANOVA was used with multiple comparisons using Sidaks or Tukey correction. Treatment data was analysed using two way ANOVA and multiple comparisons. All statistical tests were carried out using GraphPad Prism software.

## Acknowledgements

The authors would like to thank the participants from which the original skin biopsies were taken. The authors gratefully acknowledge the support of Parkinson’s UK (F-1301 awarded to HM and K-1506 awarded to HM and LF), funding from Celgene to HM, Wellcome Trust Academy of Medical Sciences Springboard funding SBF002/1142 awarded to LF and Australian Research Council (DP180101897) awarded to EN. JB and JS received funding from the SysMedPD project from the European Union’s Horizon 2020 research and innovation program under grant agreement 668738 and the ZonMW PMRare project (113302003). This research was supported by the NIHR Sheffield Biomedical Research Centre (BRC). The views expressed are those of the author(s) and not necessarily those of the NHS, the NIHR or the Department of Health and Social Care (DHSC).

## Authors contributions

AS undertook much of the experimental work culturing the neurons, performing assays and staining and putting figures together. CB undertook the remainder of neuron culture and assays. FML undertook some of the experimental work specifically the Tom20 loss mitophagy validation and KH176m mitophagy assays. MM was involved in the experimental work generating the iNPC’s from patient and control fibroblasts. EN designed and synthesised the mitochondrial ROS probes and advised on their use. JB and JS provided the Khondrion compounds and expertise of use in assays. LF was instrumental in the generation of iNPC’s both experimentally and intellectually. HM conceived the study, planned the study, undertook some experimental work, analysed and interpreted the data and wrote the manuscript. All authors contributed to the editing and re-drafting of the manuscript.

## Competing Interests

JS is the founding chief executive officer of Khondrion. JB is the chief scientific officer of Khondrion. All other authors have no competing interests to declare.

